# Legacy of past exposure to hypoxia and warming regulates ecosystem service provided by oysters

**DOI:** 10.1101/2022.04.25.488919

**Authors:** Sarah C. Donelan, Matthew B. Ogburn, Denise Breitburg

## Abstract

Climate change is having substantial impacts on organism fitness and ability to deliver critical ecosystem services, but these effects are often examined only in response to current environments. Past exposure to stress can also affect individuals via carryover effects, and whether these effects scale from individuals to influence ecosystem function and services is unclear. We explored carryover effects of two coastal climate change stressors – hypoxia and warming – on oyster (*Crassostrea virginica*) growth and nitrogen bioassimilation, an important ecosystem service. Oysters were exposed to a factorial combination of two temperature and two diel-cycling dissolved oxygen treatments at three-months-old and again one year later. Carryover effects of hypoxia and warming influenced oyster growth and nitrogen storage, with early life stress generally reducing nitrogen storage and relative tissue growth, particularly in warm environments. When extrapolated to the reef scale, carryover effects reduced estimated nitrogen storage by a restored oyster reef by as much as 41%, a substantial decline in a critical ecosystem service. Even brief exposure to climate change stressors early in life has persistent, negative effects on an ecosystem service one year later. Carryover effects on individuals impact processes at the ecological scale and must be considered in assessments of and management plans for species and ecosystems threatened by anthropogenic change.

**Significance Statement:** Anthropogenic change threatens organisms’ ability to provide ecosystem services through effects on individual phenotypes. Past experiences with anthropogenic stress can have delayed, persistent impacts on organisms via carryover effects, but how carryover effects scale to influence ecosystem function and services is not yet established. In marine systems, foundation species such as oysters mitigate effects of eutrophication by storing nutrients like nitrogen in their tissue and shell. We show that past exposure to two interacting climate change stressors (hypoxia and warming) reduces nitrogen stored by oysters by as much as 41% one year after initial exposure. Our results reveal carryover effects as a novel pathway through which climate change affects ecosystem processes that should be incorporated into conservation and management plans.

## Introduction

Climate change is altering patterns and processes across all levels of biological organization (1, 2). Phenotypic plasticity is often a key component of organismal responses to rapid environmental change (3) and can facilitate individual survival in response to changing and increasingly variable environments (e.g., 4). Plasticity in functional traits such as growth, fecundity, and foraging are critical not only for individuals and populations (5, 6), but can scale up to affect ecosystem processes such as productivity, energy flux, and nutrient cycling (7, 8). Plasticity can thereby influence species’ capacity to provide critical ecosystem services in response to changing climates and exacerbate or alleviate effects of anthropogenic change (9, 10).

Assessments of the role of plasticity in species’ responses to climate change have focused primarily on responses to current environments (e.g., 11, 12). However, organism phenotypes are also shaped by past environments such as those experienced during previous life stages. These within-generation carryover effects are widespread across taxa (13-15) and can affect important functional traits, including those that respond plastically to current environmental change and affect ecosystem functioning. For example, early life exposure to drought reduces beech tree growth and leaf production (16), which may ultimately influence their ability to sequester carbon, an important ecosystem service, as has been shown in response to current drought (17). While carryover effects are increasingly acknowledged for their impact on individuals, their connection to ecosystem processes and services has yet to be established.

In marine systems, reef building foundation species such as oysters provide physical structure that supports entire communities and are thus critical to ecosystem function (18, 19). Climate change, overfishing, and non-native species have contributed to substantial declines in many foundation species populations, threatening their ability to provide ecosystem functions and services (20-22). Oysters modify coastal biogeochemical cycles, including the nitrogen cycle, by filtering nitrogen-rich seston from the water column as they feed (23, 24). Once consumed, this nitrogen is either bioassimilated into oyster tissue and shell or excreted onto the surrounding sediment and ultimately buried or converted to nitrogen gas via microbially-mediated denitrification that then escapes from the system (24). Through these avenues, oysters provide an important ecosystem service that ameliorates downstream effects of coastal eutrophication such as algal blooms and hypoxic zones that can result in ecosystem collapse (25, 26).

While bioassimilation is only a component of this ecosystem service, it can remove nitrogen from the system for especially long periods of time (months to years, 24) and even permanently for oysters grown in aquaculture that are then harvested (27). Oysters’ ability to bioassimilate nitrogen, however, is highly plastic, varying with individual traits such as oyster size (28) and environmental factors such as low dissolved oxygen (DO, 29). DO is declining dramatically in marine systems due in part to rising atmospheric CO_2_ and temperature (30). In coastal systems, dissolved DO concentrations are affected by anthropogenic nutrient loads and seasonal changes in water temperature. They also fluctuate over shorter timescales such as day due to changes in photosynthetic rates of phytoplankton and submerged aquatic vegetation (31). The duration and magnitude of this diel-cycling hypoxia (<2 mg L^-1^, 30) are further exacerbated by warming water temperatures associated with climate change (32). Direct exposure to hypoxia and warming affects key oyster functional traits. Prolonged severe hypoxia (∼5 days) results in mass mortality of wild populations (33), and diel-cycling hypoxia can reduce oyster growth and valve gaping (necessary for feeding) by 40 and 90%, respectively (29, 34). Warm water temperatures, in contrast, generally increase oyster filtration rates (35) and growth (36). Previous work has found carryover effects of hypoxia and warming on oyster growth over short timescales: early life exposure to both hypoxia and warming resulted in a >100% reduction in relative tissue to shell growth three months later (36). But whether carryover effects persist and alter oyster bioassimilation, and hence a coastal ecosystem service, is not yet explored.

We examined carryover effects of hypoxia and warming on growth and nitrogen bioassimilation in the eastern oyster (*Crassostrea virginica*) across different environmental contexts using a multiyear laboratory experiment. Oysters were exposed to a factorial combination of two temperature (ambient, warming) and two diel-cycling DO (normoxia, diel-cycling hypoxia) treatments at three-months-old and again one year later. We measured oyster tissue and shell growth and nitrogen bioassimilated into tissue and shell during the second year to test whether carryover effects 1) influence growth and nitrogen storage in sub-adult oysters, 2) vary with subsequent exposure to the same stressors, 3) contribute to differences in the nitrogen storage potential of restored oyster reefs.

## Results

### Water quality

Complete water quality results are provided in the SI Appendix (Table S1). Briefly, DO concentrations were lower in the diel-cycling hypoxia (0.55 mg L^-1^, 7.7% saturation on average) than normoxia (7.0 mg L^-1^, 94.6% saturation on average) treatment level during the hypoxic plateau in both Year 1 and 2. Warming also reduced DO during both the normoxic and hypoxic plateaus, but these differences were very small (∼1% saturation) and likely not biologically meaningful, particularly at normoxia (34). Temperatures in the warming treatment level were 2.3°C and 2.4°C higher than ambient temperatures (Year 1 and 2, respectively). Temperature also affected pH, but differences were <0.04 units, an order of magnitude lower than levels needed to affect oyster growth (34).

### Oyster growth

Early life exposure to diel-cycling DO and warming affected oyster tissue growth regardless of Year 2 exposure (F_1,9.8_ = 7.80, p = 0.02, SI Appendix Fig. S2a), with oysters exposed to both hypoxia and warming early in life growing the same amount of tissue as control oysters (normoxia DO, ambient temperature, p = 0.07), but more tissue than oysters exposed to warming alone (8%, p = 0.01) or hypoxia alone (9%, p = 0.01) early in life. Oysters also grew 10% more tissue if they were in warm vs. ambient Year 2 temperatures (F_1,20.0_ = 5.56, p = 0.03, SI Appendix Fig. S2b). Carryover effects did not impact shell growth, but Year 2 exposure to hypoxia reduced shell growth (F_1,20.0_ = 5.35, p = 0.03, SI Appendix Fig. S3) by 8% relative to Year 2 control (p = 0.01) and 13% relative to Year 2 warm environments (p = 0.0001).

Carryover effects impacted relative tissue to shell growth and these effects varied across Year 2 environments (F_1,20.1_ = 5.04, p = 0.03, Fig. 2). In Year 2 control conditions (normoxia, ambient), oysters exposed only to warming early in life grew 14% less tissue relative to shell than oysters exposed to both warming and hypoxia early in life (p = 0.03). When oysters were exposed to multiple stressors (hypoxia and warming) in Year 2, those exposed only to hypoxia early in life tended to grow less tissue:shell than oysters exposed to both hypoxia and warming early in life (p = 0.08, 9%). Carryover effects did not impact oyster relative tissue to shell growth when oysters were exposed to only one stressor (warming or hypoxia) in Year 2.

**Figure 1.**
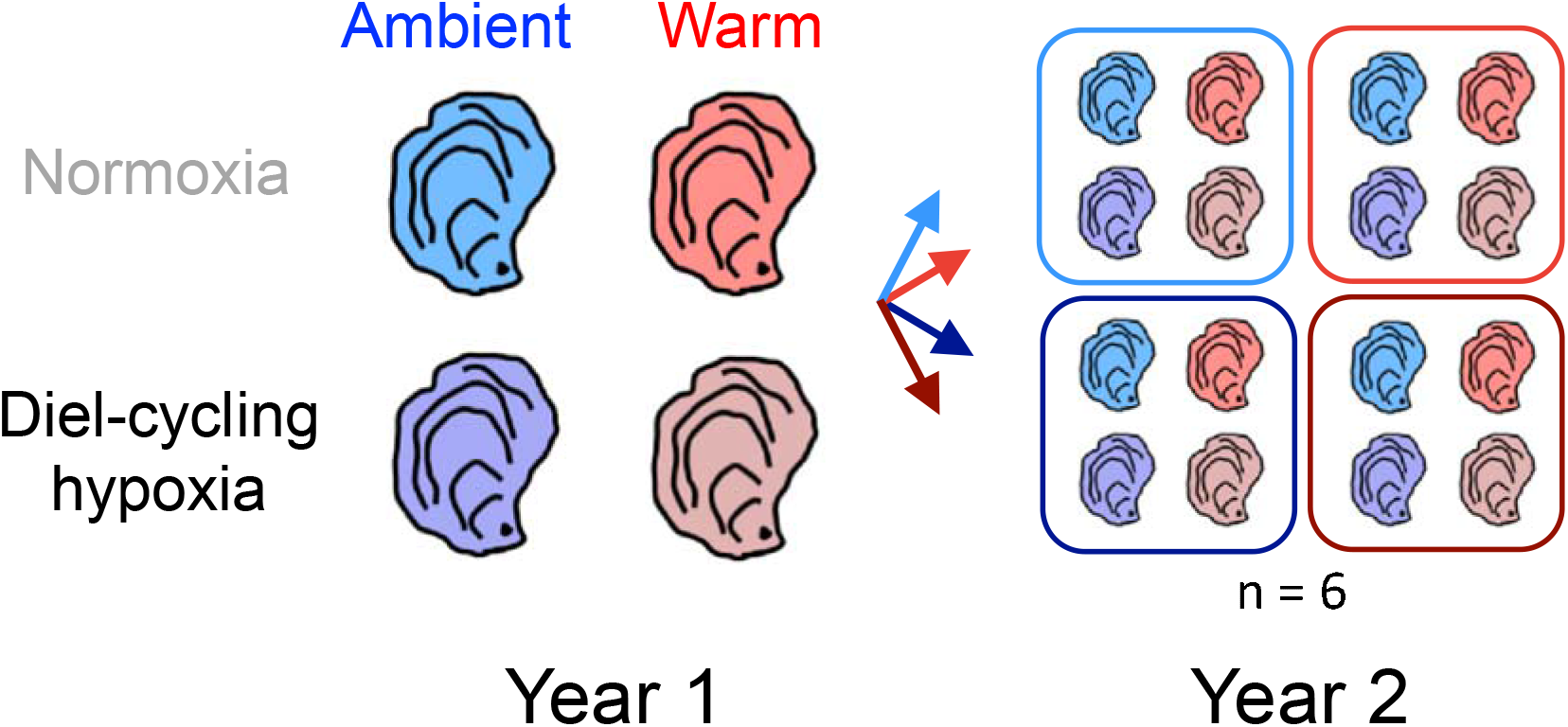
Schematic of the experimental design. Oysters (*Crassostrea virginica*) were exposed to diel-cycling dissolved oxygen (normoxia/hypoxia) and temperature (ambient/warm) treatments at three-months-old and again one year later in a fully factorial design (16 treatment combinations). See SI Appendix Table S1 for average treatment conditions.

**Figure 2.**
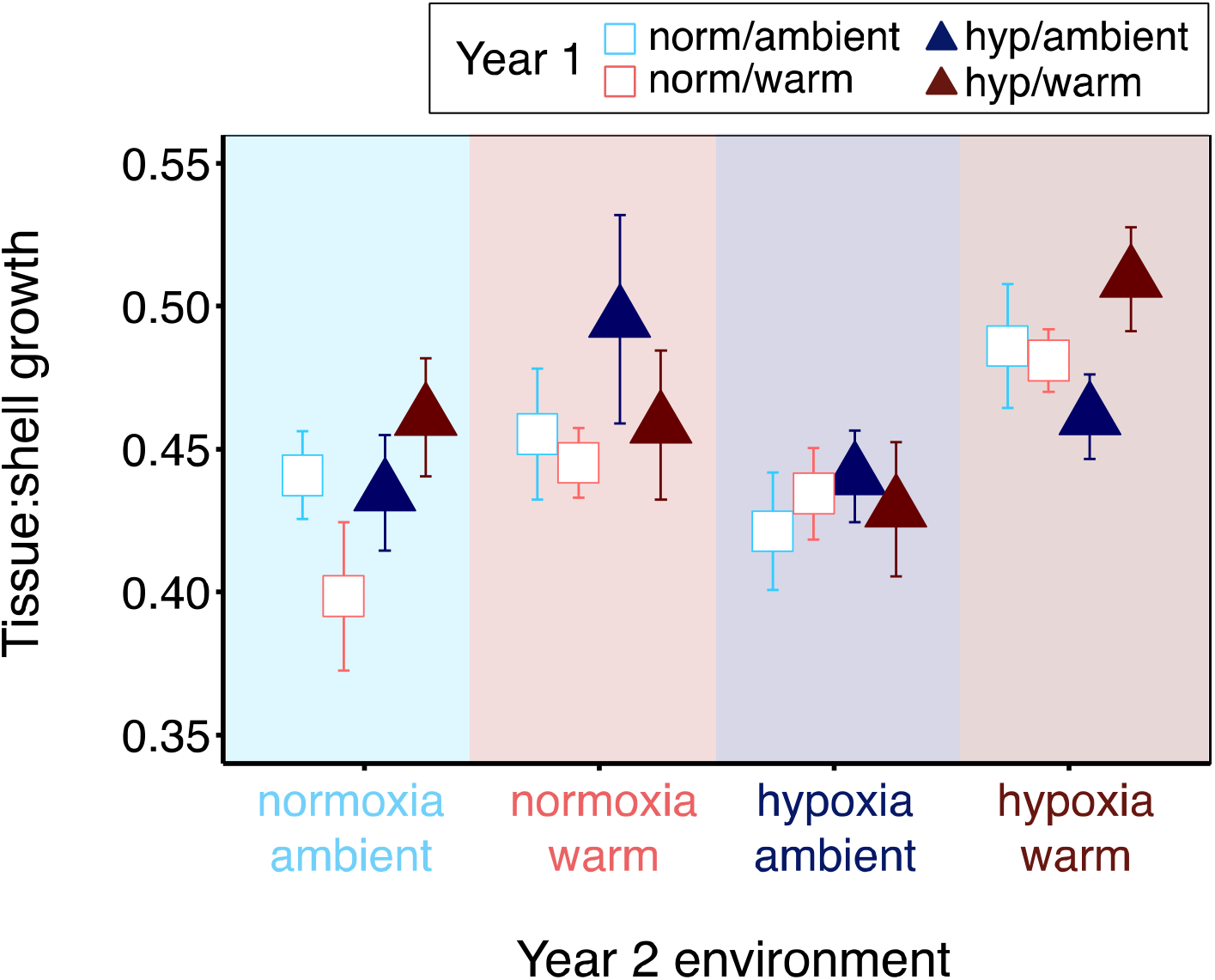
Tissue:shell growth (mean ± SEM) for oysters exposed to diel-cycling dissolved oxygen (normoxia/hypoxia) and temperature (ambient/warm) treatments at three-months-old (Year 1) and again one year later (Year 2).

### Nitrogen storage

We found carryover effects of diel-cycling DO and warming on nitrogen stored in oyster tissue and shell growth and these effects varied across Year 2 temperatures (tissue: F_1,20.1_ = 9.98, p = 0.005, Fig. 3A; shell: F_1,20.0_ = 12.34, p = 0.002, Fig. 3B). In warm Year 2 environments, oysters stored 14% less nitrogen in their tissue if they were exposed to warming alone (p = 0.01) or hypoxia alone (p = 0.004) early in life compared to oysters exposed to both hypoxia and warming or control conditions early in life (Fig. 3A). In ambient Year 2 environments, oysters stored 8% and 5% less nitrogen in their tissue growth, respectively, if they were exposed to warming alone early in life compared to oysters exposed to hypoxia alone or hypoxia and warming simultaneously (both p < 0.05). Year 2 temperature also affected nitrogen in oyster shell growth: oysters exposed to hypoxia early in life stored 14% less nitrogen in their shell than oysters exposed to normoxia early in life (p = 0.0001, Fig. 3B) and these effects were exacerbated if oysters were exposed to both hypoxia and warming early in life (18%, p = 0.0003). At ambient Year 2 temperatures, oysters exposed to both hypoxia and warming early in life had 13% less nitrogen in their shell growth compared to oysters from all other early life environments (p = 0.01).

**Figure 3.**
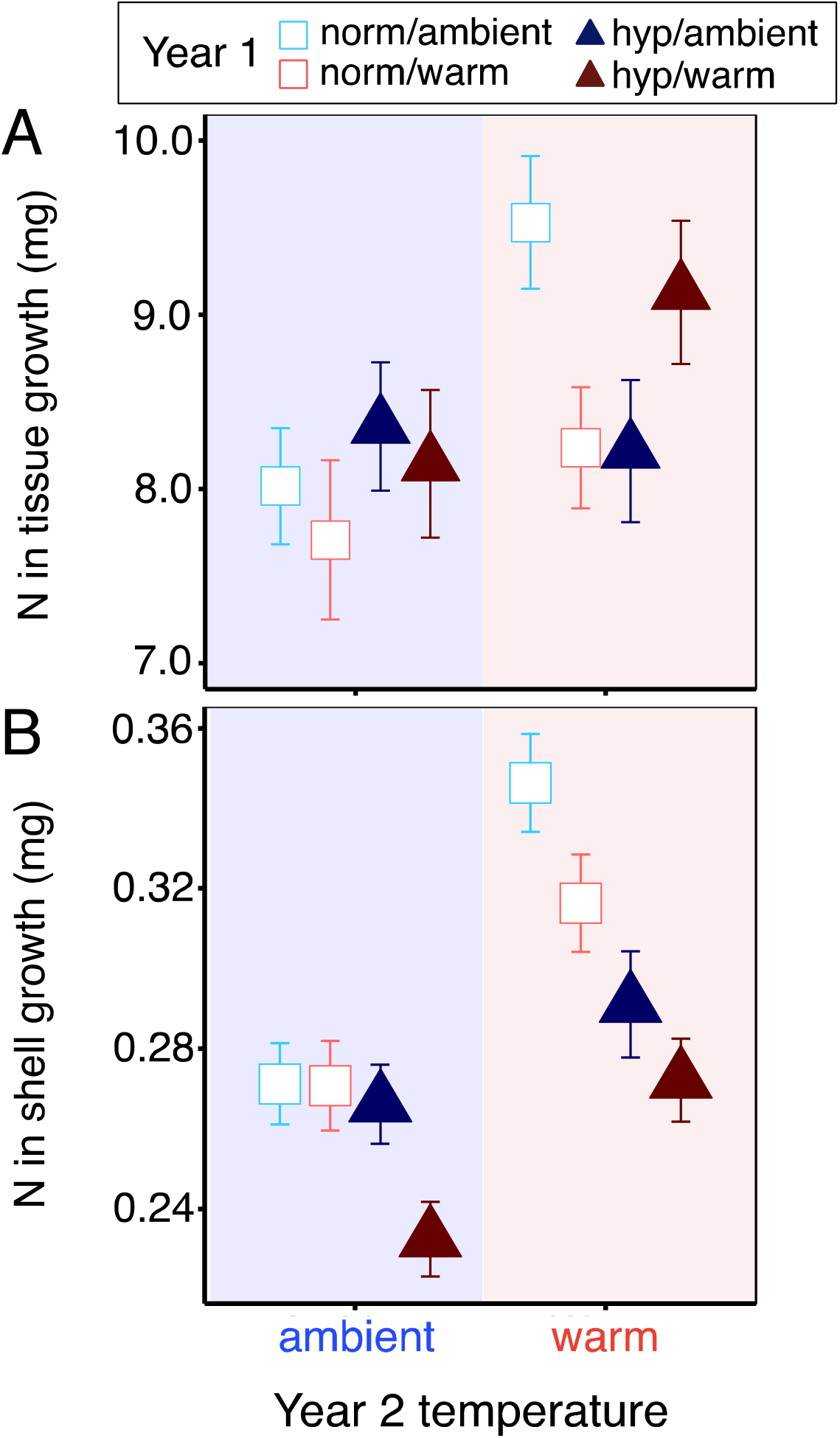
Nitrogen bioassimilated into oyster (A) tissue and (B) shell growth (mean ± SEM) at different Year 2 temperatures (x-axis) for oysters exposed to diel-cycling dissolved oxygen (normoxia/hypoxia) and temperature (ambient/warm) treatments at three-months-old (Year 1). Empirically derived percent nitrogen by weight provided for each treatment combination in SI Appendix Table S4.

Carryover effects also operated differently across Year 2 DO environments, but only for nitrogen stored in oyster shell growth (SI Appendix Table S3, Fig. S4). In normoxic Year 2 environments, early life exposure to both hypoxia and warming reduced nitrogen in oyster shell growth by 22% compared to all other early life environments (p < 0.0001). In hypoxic Year 2 environments, oysters exposed to hypoxia early in life stored 13% less nitrogen in their shell compared to oysters exposed to normoxia early in life, regardless of early life temperature (p = 0.004).

Finally, carryover effects had implications for nitrogen storage projections on a restored oyster reef, resulting in as much as a 41% reduction in N stored in an acre of reef (Fig. 4, SI Appendix Table S5). The largest decline occurred in a hypoxic potential reef environment (Fig. 4C), where carryover effects of both hypoxia and warming reduced N storage by 101 kg acre^-1^ (41%) compared to oysters from warm early life environments. Substantial declines were observed in other potential reef environments for oysters exposed to early life stress compared to early life control environments. For example, carryover effects of warming alone decreased nitrogen storage by 23% in control (67 kg acre^-1^, Fig. 4A) and warm (39 kg acre^-1^, Fig. 4B) potential reef environments, while in a hypoxic potential reef environment (Fig. 4C), early life exposure to both hypoxia and warming reduced nitrogen storage by 49 kg acre^-1^ (25%) compared to early life controls. Early life stress exposure also increased nitrogen storage in some potential reef environments; for example, in a hypoxic potential reef environment, early life exposure to warming led to a 22% increase in N stored (43 kg acre^-1^) compared to oysters from control early life conditions.

**Figure 4.**
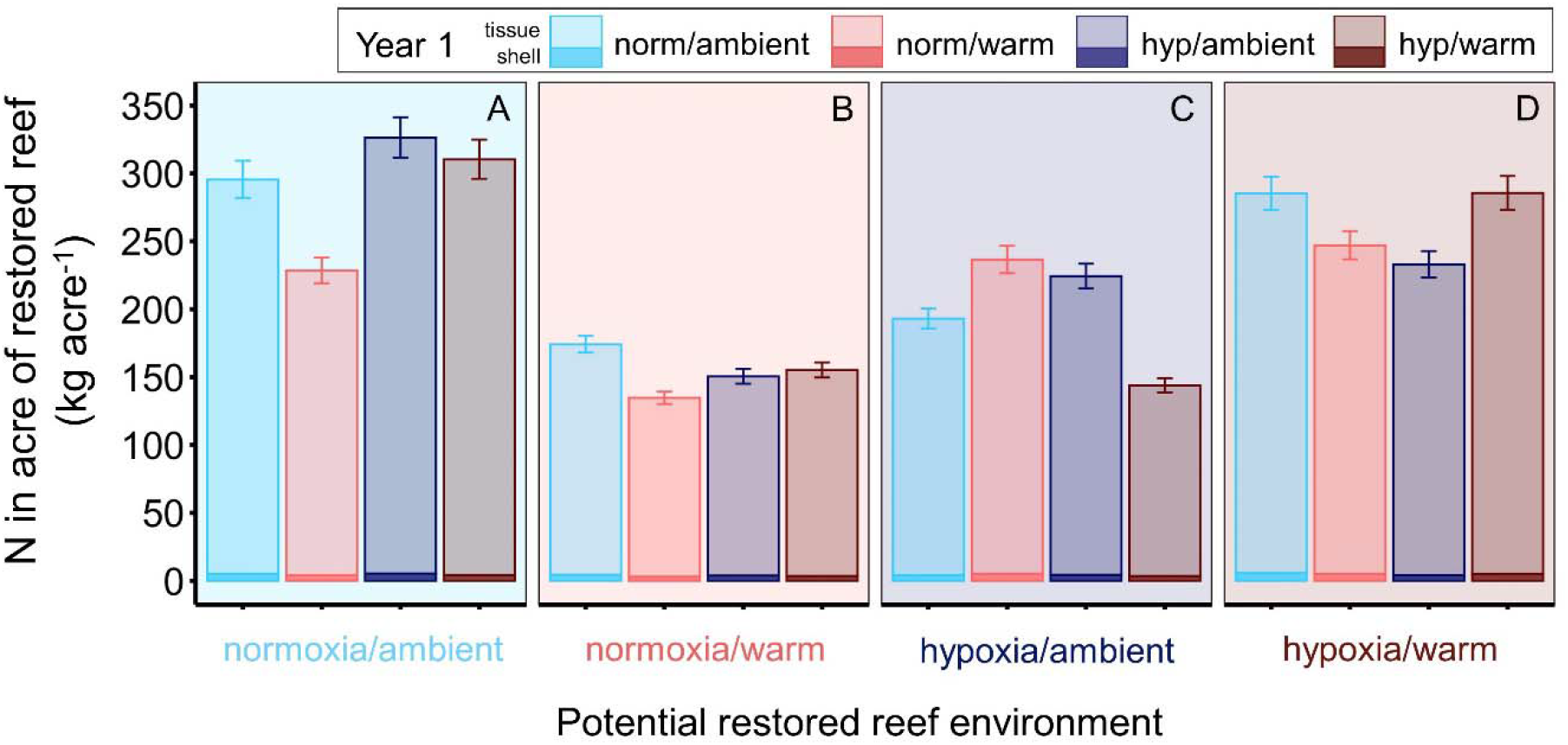
Potential nitrogen assimilated by oysters restored to Harris Creek, Maryland if oysters were exposed to different combinations of diel-cycling DO (normoxia/hypoxia) and temperature (ambient/warm) in Year 1 and outplanted to reefs that experience different hypoxia and temperature exposures. Means and 95% CI (error bars) were calculated via bootstrapping using the boot R package (43), 1000 iterations, and a normal approximation using oyster shell lengths and densities (121.42 m^-2^) taken from surveys completed in 2020 of reefs restored in 2013 (42). See SI Appendix Table S5.

## Discussion

Climate change is increasing the likelihood that organisms – particularly sessile organisms like oysters – will be exposed to multiple stressors repeatedly over their lifetimes (45). Within-generation carryover effects may therefore be increasingly common and important if they alter phenotypes in ways that are adaptive in those environments (46, 47). Moreover, changes that occur early in development are potentially more impactful than later changes because they are amplified over development and ultimately affect a greater proportion of cells in adult organisms (48). Carryover effects are indeed increasingly recognized for their ability to influence individual performance (46, 49) and population persistence (14, 50) under anthropogenic change. But how carryover effects scale from individuals to impact ecosystem function is unknown and has critical implications for estimates of ecosystem service delivery in future climates. We found that early life exposure to multiple climate change stressors has persistent carryover effects on growth, nitrogen storage, and ecosystem service delivery by an important coastal ecosystem engineer. These effects varied with environmental context, but early life stress exposure generally reduced nitrogen stored by oysters, particularly in warm environments. Our results suggest that carryover effects will have profound implications for ecosystem service delivery in a changing climate.

Carryover effects influenced nitrogen stored in oyster tissue and shell. These patterns were driven by a combination of small, non-significant changes in percent nitrogen by weight (SI Appendix Table S4) and differences in growth and are therefore reflective of variation that might exist in oysters in the wild. The majority of nitrogen (97-99%) was stored in tissue (despite being only 28% of total growth), so total nitrogen storage was dominated by these differences. Oysters generally stored more nitrogen in warm vs. ambient Year 2 environments unless they were exposed to a single stressor (hypoxia *or* warming) early in life (Fig. 3A). However, oysters exposed to multiple stressors (hypoxia *and* warming) early in life stored as much nitrogen as oysters exposed to early life control conditions (normoxia, ambient) and 14% more nitrogen than oysters exposed to either hypoxia or warming early in life. These differences were not influenced by growth alone, so are likely not driven by differences in feeding or clearance rates. Instead, early life exposure to multiple stressors may influence oyster physiology in ways that impact nitrogen assimilation efficiency (51) differently than each stressor alone. For example, early life exposure to both hypoxia and warming may favor the initiation of metabolic pathways that are nitrogen-intensive such as enhanced protein synthesis to repair damaged tissue or maintain homeostasis (52). These pathways may continue to be activated later in life even after conditions have changed because of the challenge of reversing early life phenotypes (48), thereby increasing nitrogen assimilation efficiency. Alternatively, the oyster microbiome affects denitrification rates (conversion of bioavailable nitrogen to nitrogen gas, 53) within oysters and is more taxonomically diverse in response to hypoxia (54) or warming (55). Changes in the microbiome are a putative mechanism of carryover effects in other systems (56). Differences in microbial community diversity or genetic composition driven by simultaneous exposure to hypoxia and warming early in life could permanently alter nitrogen processing rates, thereby affecting oysters’ ability to assimilate and store nitrogen over the long term.

While carryover effects of both hypoxia and warming increased nitrogen in oyster tissue growth and thus overall nitrogen storage, these oysters stored the least nitrogen in their shell growth compared to other oysters in warm Year 2 environments (Fig. 3B). Nitrogen is an important component of the proteinaceous organic matrix in shells of calcifying organisms such as oysters that is central to the biomineralization process (57). Declines in shell nitrogen content that we observed in oysters exposed to early life hypoxia (with or without warming) may therefore have important consequences for oyster shell structure and exacerbate negative effects of climate change on bivalve shell production (58). Whether nitrogen is stored in tissue or shell also has important implications for the efficacy of oysters as a mechanism for long-term nitrogen removal in coastal systems (59). Nitrogen stored in tissue is likely removed for a shorter time than nitrogen in shell because oyster shells maintain their integrity even after death while tissue readily decomposes, releasing nitrogen back into the system (24).

Carryover effects also operated differently across Year 2 DO environments, but only through effects on nitrogen in oyster shell, not tissue, growth. Because shell nitrogen storage accounts for only 1-3% of overall nitrogen stored, these changes were small compared to differences in carryover effects between Year 2 warming environments. Our results therefore suggest that carryover effects will be more impactful in warm versus hypoxic environments, at least for nitrogen storage. DO concentrations in the environment may not have strong effects on nitrogen assimilation within oysters themselves, perhaps due to changes in the microbiome associated with early life hypoxia or oysters’ ability to avoid exposure to hypoxia by reducing feeding during hypoxic periods (29).

When extrapolated to the reef scale, carryover effects have critical implications for the potential of oysters to deliver ecosystem services. Differences in early life environments reduced estimated reef storage capacity by as much as 41% (101 kg acre^-1^, Fig. 4C). Compared to reefs comprised of oysters exposed to control conditions early in life, reefs of oysters exposed to single or multiple stressors early in life are predicted to store up to 25% less nitrogen (67 kg acre^-1^). Harris Creek is one of the largest oyster restoration projects in the world, with 348 acres of reef restored (42, 60). On this reef scale, carryover effects have the potential to reduce nitrogen stored long-term in oyster tissue and shell by ∼35,100 kg (35.1 metric tons), a profound change in ecosystem service potential. In-water removal of nitrogen by bivalves is often cited as a key benefit of habitat restoration projects and aquaculture (27) and increasingly considered in watershed management plans enacted to improve water quality (61). For aquaculture operations, there is also interest in compensating growers for nutrients removed via their leases through nutrient trading programs (27, 62). But estimates of nitrogen bioassimilated by oysters are often based only on oyster size (59) and/or periodic measurements (e.g., monthly) of relevant environmental parameters (27). Our results show that carryover effects can considerably limit the capacity of oysters to deliver this service and thus are critical to incorporate into estimates of nutrient reduction provided by oyster restoration and aquaculture. Indeed, the 101 kg acre^-1^ change we observed as a result of carryover effects is similar to nitrogen stored by oysters in an acre of aquaculture farm over an entire year in Chesapeake Bay (27).

We also found context-dependent carryover effects on oyster relative tissue to shell growth. Oysters exposed to hypoxic, warm environments early in life grew more tissue relative to shell than oysters exposed to a single stressor early in life in multiple Year 2 environments (control and hypoxia, warm, Fig. 2). These results oppose those of our earlier work that found negative effects of early life exposure to both hypoxia and warming on relative tissue growth, particularly when oysters were re-exposed to those conditions three months later (36). These contrasting patterns emphasize the need to quantify carryover effects across multiple stages of ontogeny. Because tissue mass is a proxy for oyster fitness (63), carryover effects that impact tissue growth may be increasingly influential as oysters approach sexual maturity (∼30 mm shell length, 63). At ∼14 mm shell length on average, our experimental oysters were small for their age, likely due to abnormally low salinity in the Chesapeake Bay across experimental years (36, 64), and still sexually immature. But if the positive trends we observed here continue to influence oysters as they approach sexual maturity, early life exposure to multiple climate change stressors may actually increase oyster fecundity and alter oyster sex ratios (oysters are protandrous hermaphrodites, 63) in particular environments. These patterns should be explicitly explored in future work.

Our results reveal that early life exposure to multiple climate change stressors has context-dependent effects on oyster growth and nitrogen storage and implications for the ability of oysters to mitigate eutrophication in coastal systems. Not only were carryover effects persistent, affecting oysters one year after initial exposure, but they varied across the life cycle (36), suggesting that impacts on nitrogen removal may further vary by life stage or time since initial exposure to stress. Our research demonstrates the need to incorporate carryover effects not only into assessments of individual and population performance as others have done (e.g., 49, 50), but also into assessments of ecosystem function and service provisioning. Such steps will be challenging, but critical, to accurately assess the impact of climate change in natural systems.

## Materials and Methods

### Experimental design and set-up

We conducted a multiyear fully factorial experiment to explore how early life exposure (Year 1) to diel-cycling dissolved oxygen (DO, normoxia/hypoxia) and temperature (ambient/warm) affected oyster tissue, shell, and relative tissue:shell growth and total nitrogen bioassimilated in tissue and shell growth when oysters were exposed to diel-cycling DO (normoxia/hypoxia) and temperature (ambient/warm) treatments one year after initial exposure (Year 2). There were a total of 16 treatment combinations (Fig. 1). The Year 1 exposure ran in August 2018 and the Year 2 exposure in August 2019. Experimental methods presented here are similar to those in Donelan, Breitburg and Ogburn (36) and all relevant details not described here are provided there and in the SI Appendix.

Briefly, 3-4 month old oysters were purchased from Horn Point Oyster Hatchery (Cambridge, Maryland) and acclimated for six days in flow-through aquarium facilities at the Smithsonian Environmental Research Center (SERC) on the Rhode River, Maryland prior to the Year 1 exposure. Conditions were similar between the two locations (within 2 psu salinity and 1.5°C), so oysters should not have been physiologically stressed by this change (37).

The experiment took place in an indoor aquarium facility at SERC that manipulates the temperature and DO concentration of untreated Rhode River water. Water in the warming treatment level was dynamically maintained at 2.5°C above ambient to match predictions of the 2014 IPCC RCP6.0 scenario (38). Water temperature was unmanipulated in the ambient treatment level. DO concentrations in the normoxic treatment level were held at normoxia (∼100% saturation). DO concentrations in the diel-cycling hypoxia treatment level were manipulated over a 24-hour cycle using custom LabView software (detailed in 36, 39) that uses a mixture of gases and continuous monitoring of one tank from each treatment combination to achieve desired concentrations (details in SI Appendix). A hypoxic cycle consisted of a 3-hour draw down from normoxia to hypoxia (target: 0.5 mg L^-1^, 7-8% saturation), a 4-hour plateau period when hypoxia was maintained, a 3-hour ramp up from hypoxia to normoxia, and a 14-hour period at normoxia, mimicking DO concentrations and cycle durations in tributaries of the Chesapeake Bay (40). pH remained constant by adding CO_2_ gas as needed and was similar to ambient conditions during the experiment. Each aquarium was covered with a tight-fitting plexiglass lid to maintain target DO, temperature, and pH, which were verified in each tank once or twice during a daily hypoxic plateau using an external water quality probe. DO cycled five days each week in the diel-cycling treatment and stayed at normoxia on the remaining two days, except for one week in Year 2 when it cycled for seven days (SI Appendix, Figure S1). This schedule was used for logistical reasons, but also mirrors natural variability (40). The temperature treatment was applied each day, again mimicking natural patterns.

### Growth and nitrogen storage

At the start of the Year 1 exposure, oysters (N = 3,000) were placed into the experimental array. There were six replicate tanks of each of the four treatment combinations (N = 24, n = 150 oysters tank^-1^). In each tank, oysters were held in a perforated plastic container (16 × 16 × 9.5 cm, l × w × h) that was placed on a semi-open platform to minimize sedimentation. Oysters remained in these conditions for 18 days (13 days of diel-cycling DO, 18 days of temperature treatment). At the end of the Year 1 exposure, oysters were placed in a tank of aerated, ambient Rhode River water for two months before being moved (within their experimental mesocosms) to large vexar mesh bags (84 × 48 × 10 cm, l × w × h; 9 mm mesh) suspended off the SERC dock in the Rhode River until the start of the Year 2 exposure. Bags were rinsed throughout the winter to prevent sedimentation, fouling, and oyster mortality.

At the start of the Year 2 exposure, we placed approximately half of the oysters (N = 960) from Year 1 back into the experimental array. We standardized oyster size (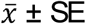; shell mass: 412.8 ± 5.1 mg, p = 0.9, tissue mass: 176.1 ± 2.7 mg, p = 0.3) to remove potential effects on growth patterns. We did this by excluding oysters from one replicate of each Year 1 treatment combination that on average contained the largest oysters. Hence, oysters from five replicates of each Year 1 treatment combination were used in Year 2. There were 24 tanks with 10 oysters from each Year 1 treatment combination in each tank (n = 40 oysters tank^-1^, 1-3 oysters Year 1 replicate^-1^). Oysters remained in the Year 2 exposure for 20 days (16 days of diel-cycling DO, 20 days of temperature treatment).

To track individual oyster tissue and shell growth, we tagged each oyster with a plastic numeric tag and weighed them using the buoyant weighing technique (36, 41) immediately prior to the start and following the end of the Year 2 exposure (details in SI Appendix). Shell and tissue growth (mg) were calculated as final – initial mass. Relative tissue:shell growth was calculated by dividing individual tissue growth by shell growth.

Immediately after weighing, a subsample of oysters (n = 6, N = 96) were placed in a -80°C freezer. Nine months later, they were removed, dissected to separate tissue and shell, and oven dried at 60°C. Each shell and tissue sample was then ground into a fine powder using a ball mill grinder, weighed, packed individually into tin capsules, and analyzed for the percent of total (organic + inorganic) nitrogen by weight on a Thermo Delta V Advantage mass spectrometer coupled to an Elemental vario ISOTOPE Cube Elemental Analyzer at the Smithsonian MCI Stable Isotope Mass Spectrometry Laboratory. This generated an average percent nitrogen in oyster tissue and shell for each treatment combination (SI Appendix Table S4), which was multiplied by the tissue and shell growth of each oyster as appropriate to determine the nitrogen contained in tissue and shell growth (mg).

### Nitrogen storage projections for a restored reef

We used the nitrogen content of the experimental oysters to estimate how carryover effects may impact nitrogen stored on a restored oyster reef. We used a restored reef in Harris Creek, a tributary of the Choptank River in Maryland, as a model because it was restored with oysters from the same population as our experimental oysters, is one of the oldest, most successful oyster restoration projects in Maryland, and has been regularly monitored for oyster size, density, and spatial extent. Oyster size and density data from Harris Creek were obtained from the 2020 Maryland Oyster Monitoring Report using reefs restored with live oysters in 2013 and monitored for shell length and density in 2020 (42). Harris Creek oysters were substantially larger (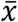 shell length ± SE; 73.6 ± 0.62 mm) than the experimental oysters (13.6 ± 0.09 mm), but the experimental oysters also differed in size across treatment combination. To capture these size differences in potential effects on nitrogen storage, we used allometric scaling to extrapolate the tissue and shell mass of an oyster in Harris Creek based on relationships between shell length (mm) and tissue and shell mass (mg) of the experimental oysters. Specifically, we used log-log plots to generate relationships between shell length and tissue and shell mass (separately) for oysters from each of the 16 experimental treatment combinations (SI Appendix Table S5). We used these equations to bootstrap (1000 iterations, boot package, 43) mean and 95% confidence intervals of nitrogen stored in oyster tissue and shell in a square meter of restored reef using the shell lengths and density from the 2020 report (121.42 oysters m^-2^, 42).

### Statistical analyses

DO concentrations, pH, and temperature were analyzed to determine differences between treatments. Year 1 and Year 2 were analyzed separately. Complete analyses are described in the SI Appendix. We analyzed oyster tissue growth (mg), shell growth (mg), tissue:shell growth, and nitrogen stored in oyster tissue and shell growth (mg) using separate linear mixed models with Year 1 DO, Year 1 temperature, Year 2 DO, and Year 2 temperature as fixed effects and REML variance estimation. Initial shell mass (mg) was included as a covariate and significant for all analyses (p < 0.04, Table S1). To account for non-independence among oysters within the same tank, the following nested factors were included as random effects: Year 1 replicate nested within Year 1 DO and Year 1 temperature, Year 2 replicate nested within Year 2 DO and Year 2 replicate, and Year 2 replicate was separately crossed with Year 1 DO, Year 1 temp, and Year 1 DO x Year 1 Temp and nested within Year 2 DO and Year 2 temperature. F-ratios and p-values were generated using Type III ANCOVAs (SI Appendix Tables S2, S3). Data met the assumptions of ANCOVA except the residuals were not normal, but ANCOVA is robust to violations of normality. Post hocs were calculated using ls means (α = 0.05). Analyses were conducted in R (v. 3.6.3, 44). Three oysters died during the experiment, so were excluded from analyses.

## Acknowledgements

We thank Kristina Borst, Catherine Bubser, Christine France, Daniella Gavriel, and Sarah Gignoux-Wolfsohn for help with the experiment and sample processing, Roy Rich for lending the temperature manipulation equipment, and Shannon Hood and Alex Golding at Horn Point Oyster Hatchery for assistance with oyster spat. This work was supported by a Smithsonian Institution Postdoctoral Fellowship to S.C. Donelan, the National Science Foundation (DBI-1659668), and Maryland Sea Grant Award #NA14OAR4170090-SA75281450-P to D. Breitburg and M.B. Ogburn.

