## Supplementary material for "Legacy of past exposure to hypoxia and warming regulates ecosystem service provided by oysters": SI Appendix

SI Appendix for

### Methods

*Experimental oyster origin.* Experimental oysters were spawned at Horn Point Oyster Hatchery in Cambridge, Maryland from broodstock ($n_{female}$ = 39, $n_{male}$ = 16) collected from the Choptank River, a tributary of the Chesapeake Bay. We purchased the oysters when they were 3-4 months old (3-5 mm shell height) and moved them to the flow-through aquarium facilities at the Smithsonian Environmental Research Center (SERC) on the Rhode River, another tributary of the Chesapeake Bay, in Edgewater, Maryland.

*Experimental set-up and treatment manipulations.* The experiment took place in an indoor aquarium facility at SERC that pumped untreated water from the Rhode River into a 568L fiberglass holding tank set in series with four other identical holding tanks. The first three tanks allowed sediment to settle and water was pumped from the final two tanks into experimental aquaria. One tank that fed the experimental aquaria remained at ambient temperatures while one was warmed to a target water temperature of 2.5°C above ambient. Warming was dynamically maintained through a custom microprocessor feedback control (1) that controlled power to 3000 watts of aquarium heaters (500W heaters, Aquatop, Brea, California, USA). Water was dispensed from the ambient and warm holding tanks into 75L experimental aquaria through vinyl tubing at a rate of 300 mL $min^{-1}$.

DO concentrations were controlled with custom LabView software (2) that uses input from dissolved oxygen (Oxyguard, Birkeroed, Denmark) and pH (Durafet III, Honeywell, Fort Washington, Pennsylvania, USA) sensors placed in one replicate tank of each treatment combination. The program then controls the flow of a mixture of four gases (nitrogen, carbon dioxide, air, and $CO_2$-stripped air) as necessary to achieve target DO concentrations for a given timepoint. The program also maintains constant pH by adding $CO_2$ as needed, and pH was similar to ambient conditions in the Rhode River during each year of the experiment (target pH: Year 1: 8.1, Year 2: 8.0, Table S1). Gases were dispensed to each replicate aquarium at a rate of 25 L $min^{-1}$ through mass flow controllers (Dakota Instruments, Orangeburg, New York, USA), gas manifolds, vinyl tubing, and silica airstones. Each aquarium was covered with a tight-fitting plexiglass lid to minimize ambient gas exchange.

The four continuously monitoring DO and pH sensors that provided input to the software were checked and calibrated on almost all mornings using an external probe (Orion Star A326, Thermo Scientific, Waltham, Massachusetts, USA). This external probe reading was used in our water quality analysis as the daily DO reading ($n_{Year1} = 11$, $n_{Year2} = 10$). The same external probe was used to measure DO concentration, temperature, and pH in all 24 replicate tanks once or twice during the daily hypoxic plateau ($n_{Year1} = 26$, $n_{Year2} = 20$) to verify that the targets were reached. Readings taken during a hypoxic plateau were used in our water quality analysis as the hypoxic DO and temperature readings. We used pH readings from both the normoxic and hypoxic tank readings ($n_{Year1} = 37$, $n_{Year2} = 30$) in our water quality analysis. We also measured alkalinity in one replicate of the normoxic, ambient and normoxic, warm treatment combinations ($n_{Year1} = 3$, $n_{Year2} = 5$) and used this to calculate $pCO_2$ (uatm) using the seacarb package in R (3).

Lights were kept on a 14 h light:10 h dark cycle to simulate conditions at ~2 m depth in the Rhode River where an oyster reef might be found (4).

*Buoyant weighing technique.* Oysters were placed in a basket that was hung from the underside of a balance (Mettler Toledo AE 163) and was submerged in room temperature, Rhode River water to measure approximate oyster shell mass. Oysters were then placed on a paper towel to dry at room temperature for 1 hour, after which time they were weighed in air to measure their total dry mass (tissue + shell). Using an empirically derived equation (actual shell mass = 1.4315*submerged mass + 0.0188; $R^2 = 0.9875$) obtained through a destructive regression (5), we calculated actual oyster shell mass (mg) from the approximate oyster shell mass obtained through buoyant weighing. We calculated oyster tissue mass (mg) by subtracting actual shell mass from total dry mass sensu (5). Oysters were measured immediately before and after the Year 2 exposure and growth calculated as final – initial mass.

*Statistical analyses*

*Water quality.* We analyzed dissolved oxygen (DO) concentration during the normoxic plateau, DO concentration during the hypoxic plateau, temperature, and pH to measure differences between our experimental treatments (measured as described above) using two-way ANOVAs that considered diel-cycling dissolved oxygen and temperature as fixed effects. We also included replicate nested within diel-cycling DO and temperature as appropriate to account for potential non-independence among readings and used REML variance estimates. Alkalinity and pCO2 were analyzed using a one-way ANOVA that considered temperature as a fixed effect.

### Results

*Water quality*. During the hypoxic plateau, DO concentrations were significantly lower in the diel-cycling hypoxia than normoxia treatment level (Year 1 DO: $F_{1,16} = 75{,}170.6$, $p < 0.0001$;

Year 2 DO: $F_{1,20} = 174,133.6$, $p < 0.0001$) and very closely matched our target hypoxia value (Table S1, 0.5 mg $L^{-1}$, 7-8% saturation). During the normoxic plateau, warming treatments had significantly lower DO concentrations than ambient temperature treatments (Year 1 Temp: $F_{1,40} = 25.1$, $p < 0.0001$; Year 2 Temp: $F_{1,36} = 34.7$, $p < 0.0001$; Table S1) regardless of diel-cycling DO (both Year 1 DO and Year 2 DO $p > 0.6$). This effect of temperature on DO concentration persisted during the hypoxic plateau: in Year 1, warming reduced DO only in the normoxia treatment (Year 1 DO x Year 1 Temp: $F_{1,16} = 76.5$, $p < 0.0001$; Table S1). In Year 2, DO in the normoxia, warming and hypoxia, warming treatments were lower than their respective ambient temperature treatment levels (Year 2 DO x Year 2 Temp: $F_{1,20} = 88.3$, $p < 0.0001$; Table S1). While statistically significant, these differences in dissolved oxygen were very small (~1% saturation) and therefore likely not biologically meaningful, particularly at high oxygen levels (6).

Temperatures in the warming treatments were significantly higher than those in the ambient treatment level (Year 1 Temp: $F_{1,16} = 1087.2$, $p < 0.0001$, Year 2 Temp: $F_{1,20} = 320.8$, $p < 0.0001$, Table S1). Warm temperature treatment levels were 2.3°C and 2.4°C higher than ambient temperatures in Year 1 and Year 2, respectively, and closely matched our target temperature difference (2.5°C). Oysters also frequently experience temperatures within this range (7).

Temperature also affected pH in Year 1 ($F_{1,16.0} = 33.4$, $p < 0.0001$) and Year 2, with lower pH in the warm versus ambient temperature treatment levels (Table S1), though this pattern only emerged in the diel-cycling DO treatment in Year 2 (Year 2 DO x Year 2 Temp: $F_{1,20.0} = 15.8$, $p = 0.0007$; Table S1). However, these pH differences were <0.04 unit range, which is an order of magnitude lower than levels needed to elicit changes in growth in this system (6, 8). Finally, alkalinity and $pCO_2$ were similar across all treatments in Year 1 (alkalinity: $F_{1,4} = 7.3$, $p = 0.06$, $pCO_2$: $F_{1,4} = 0.57$, $p = 0.49$) and Year 2 (alkalinity: $F_{1,8} = 0.05$, $p = 0.83$, $pCO_2$: $F_{1,8} = 0.31$, $p = 0.59$, Table S1).

**Table S1.** Water quality measurements (mean ± SEM) during Year 1 (August 2018) and Year 2 (August 2019). Dissolved oxygen (DO) was measured by one method during the normoxic phase and another method during the hypoxic phase of a diel-cycle in a given day: in the normoxic phase, data were taken from one replicate of each treatment combination once daily on most days using an external probe (Thermo Scientific Orion Star A326, $n_{Year1} = 11$, $n_{Year2} = 10$); in the hypoxic phase, data were taken once or twice during the hypoxic plateau in each replicate tank with the same external probe ($n_{Year1} = 26$, $n_{Year2} = 20$). Dissolved oxygen was measured in mg $L^{-1}$, but we also report average % saturation. Temperature (°C) was measured once or twice daily in each tank at the same time as the hypoxic DO readings ($n_{Year1} = 26$, $n_{Year2} = 20$), and pH was measured two or three times daily at the same time as the normoxic and hypoxic DO readings (both with the same external probe). Salinity (practical salinity units, psu) readings were taken with an EXO2 sonde (YSI, Yellow Springs, Ohio, USA) mounted in the Rhode River at the same depth (~2 m) as the intake pipe for the flowing water system that recorded conditions every 6 minutes ($n_{Year1} = 4{,}079$, $n_{Year2} = 4{,}800$). Finally, water samples were taken for alkalinity (umoles/kg seawater) three times in Year 1 and five times in Year 2 and processed according to the methods described in (9). Letters indicate significant differences based on ANOVAs (see Results).

**Table S1.**

| | | *Treatments* | | | |
|---|---|---|---|---|---|
| | | Normoxia | | Diel-cycling hypoxia | |
| | | Ambient | Warm | Ambient | Warm |
| **Year 1 exposure** | | | | | |
| Dissolved oxygen (mg $L^{-1}$) | Normoxia | 6.77 ± 0.05$^{A}$ | 6.51 ± 0.04$^{B}$ | 6.75 ± 0.06$^{A}$ | 6.56 ± 0.03$^{B}$ |
| | Hypoxic plateau | 6.93 ± 0.02$^{A}$ | 6.51 ± 0.01$^{B}$ | 0.56 ± 0.02$^{C}$ | 0.55 ± 0.03$^{C}$ |
| Dissolved oxygen (% sat) | Normoxia | 88.8 ± 0.64$^{A}$ | 89.0 ± 0.57$^{B}$ | 88.6 ± 0.74$^{A}$ | 89.6 ± 0.43$^{B}$ |
| | Hypoxic plateau | 91.1 ± 0.18$^{A}$ | 89.2 ± 0.14$^{B}$ | 7.33 ± 0.24$^{C}$ | 7.22 ± 0.07$^{C}$ |
| Temperature (°C) | | 28.30 ± 0.07$^{A}$ | 30.68 ± 0.07$^{B}$ | 28.27 ± 0.08$^{A}$ | 30.61 ±0.07$^{B}$ |
| pH | | 8.13 ± 0.002$^{A}$ | 8.10 ± 0.002$^{B}$ | 8.14 ± 0.002$^{A}$ | 8.11 ± 0.002$^{B}$ |
| Salinity (psu) | | 4.00 ± 0.13 | 4.00 ± 0.13 | 4.00 ± 0.13 | 4.00 ± 0.13 |
| Alkalinity (umol/kg sw) | | 1315.3 ± 3.1$^{A}$ | 1305.3 ± 5.6$^{A}$ | | |
| $pCO_2$ (uatm) | | 360.7 ± 6.4$^{A}$ | 376.8 ± 20.5$^{A}$ | | |
| **Year 2 exposure** | | | | | |
| Dissolved oxygen (mg $L^{-1}$) | Normoxia | 7.23 ± 0.07$^{A}$ | 6.84 ± 0.06$^{B}$ | 7.25 ± 0.06$^{A}$ | 6.89 ± 0.06$^{B}$ |
| | Hypoxic plateau | 7.49 ± 0.02$^{A}$ | 7.08 ± 0.02$^{B}$ | 0.60 ± 0.006$^{C}$ | 0.50 ± 0.008$^{D}$ |
| Dissolved oxygen (% sat) | Normoxia | 96.4 ± 0.9$^{A}$ | 95.0 ± 0.8$^{B}$ | 96.5 ± 0.8$^{A}$ | 95.7 ± 0.8$^{B}$ |
| | Hypoxic plateau | 99.8 ± 0.2$^{A,C}$ | 98.4 ± 0.19$^{A,D}$ | 8.05 ± 0.08$^{B,C}$ | 6.93 ± 0.11$^{B,D}$ |
| Temperature (°C) | | 27.81 ± 0.14$^{A}$ | 30.26 ± 0.13$^{B}$ | 27.76 ± 0.14$^{A}$ | 30.19 ± 0.14$^{B}$ |
| pH | | 8.01 ± 0.003$^{AB}$ | 7.99 ± 0.004$^{B}$ | 8.00 ± 0.004$^{B}$ | 8.03 ± 0.005$^{A}$ |
| Salinity (psu) | | 8.17 ± 0.76 | 8.17 ± 0.76 | 8.17 ± 0.76 | 8.17 ± 0.76 |
| Alkalinity (umol/kg sw) | | 1415.6 ± 31.2$^{A}$ | 1425.8 ± 34.7$^{A}$ | | |
| $pCO_2$ (uatm) | | 472.3± 7.86$^{A}$ | 478.0± 6.58$^{A}$ | | |

**Table S2.** Summary of results from linear effects models and ANCOVAs that examined the effects of Year 1 diel-cycling dissolved oxygen (Year 1 DO), Year 1 temperature (Year 1 Temp), Year 2 diel-cycling dissolved oxygen (Year 2 DO), and Year 2 temperature (Year 2 Temp) on oyster a) tissue, b) shell, and c) tissue:shell growth (mg). Initial shell mass (mg) was used as a covariate. We also included the following nested terms as random effects to account for the nested design and non-independence among individuals in Year 1 and 2: Year 1 replicate was nested within Year 1 DO and Year 1 Temp, Year 2 replicate was nested within Year 2 DO and Year 2 Temp, and Year 2 replicate was separately crossed with Year 1 DO, Year 1 Temp, and Year 1 DO x Year 1 Temp and nested within Year 2 DO and Year 2 Temp. Analyses were conducted in R v. 3.6.3 (10) using the lme4 (11), car (12), and emmeans (13) packages.

| | a) tissue growth (mg) | | b) shell growth (mg) | | c) tissue:shell growth | |
|---|---|---|---|---|---|---|
| **Effect** | **F $(df_{num}, df_{den})$** | **p** | **F $(df_{num}, df_{den})$** | **p** | **F $(df_{num}, df_{den})$** | **p** |
| Year 1 DO | 1.91 (1,15.6) | 0.19 | 2.29 (1,16.9) | 0.15 | 2.00 (1,11.5) | 0.18 |
| Year 1 Temp | 2.25 (1,11.1) | 0.16 | 3.40 (1,12.5) | 0.09 | 0.008 (1,9.5) | 0.93 |
| Year 2 DO | 2.60 (1,20.0) | 0.12 | 18.51 (1,20.0) | **0.0003** | 0.37 (1,20.0) | 0.55 |
| Year 2 Temp | 5.56 (1,20.0) | **0.03** | 0.14 (1,20.0) | 0.71 | 7.69 (1,20.0) | **0.01** |
| Y1 DO*Y1 Temp | 7.80 (1,9.8) | **0.02** | 3.47 (1,10.0) | 0.09 | 0.60 (1,9.5) | 0.46 |
| Y1 DO*Y2 DO | 1.51 (1,20.0) | 0.23 | 0.51 (1,19.9) | 0.49 | 1.31 (1,19.9) | 0.26 |
| Y1 DO*Y2 Temp | 0.09 (1,20.0) | 0.77 | 0.06 (1,19.9) | 0.81 | 0.04 (1,19.8) | 0.84 |
| Y1 Temp*Y2 DO | 1.67 (1,19.9) | 0.21 | 1.21 (1,19.7) | 0.29 | 2.14 (1,19.9) | 0.16 |
| Y1 Temp*Y2 Temp | 0.41 (1,19.9) | 0.53 | 4.63 (1,19.7) | **0.04** | 0.04 (1,20.0) | 0.85 |
| Y2 DO*Y2 Temp | 1.04 (1,20.0) | 0.32 | 5.35 (1,20.0) | **0.03** | 0.61 (1,20.0) | 0.44 |
| Y1 Temp*Y1 DO*Y2 DO | 1.78 (1,20.2) | 0.20 | 0.70 (1,20.2) | 0.41 | 0.06 (1,20.2) | 0.80 |
| Y1 Temp*Y1 DO*Y2 Temp | 0.07 (1,20.0) | 0.80 | 0.30 (1,20.1) | 0.59 | 0.05 (1,20.1) | 0.83 |
| Y2 Temp*Y1 DO*Y2 DO | 0.02 (1,20.0) | 0.88 | 0.005 (1,19.9) | 0.94 | 0.009 (1,19.8) | 0.93 |
| Y2 Temp*Y1 Temp*Y2 DO | 0.41 (1,19.9) | 0.53 | 1.19 (1,19.7) | 0.29 | 1.01 (1,20.0) | 0.33 |
| Y1 Temp*Y1 DO*Y2 Temp*Y2 DO | 1.36 (1,20.0) | 0.26 | 0.65 (1,20.1) | 0.43 | 5.04 (1,20.1) | **0.03** |
| Initial Shell Mass (mg) | 727.9 (1,924.8) | **<0.0001** | 1961.7 (1,930.4) | **<0.0001** | 4.15 (1, 918.9) | **0.04** |

**Table S3.** Summary of results from linear effects models and ANCOVAs that examined the effects of Year 1 diel-cycling dissolved oxygen (Year 1 DO), Year 1 temperature (Year 1 Temp), Year 2 diel-cycling dissolved oxygen (Year 2 DO), and Year 2 temperature (Year 2 Temp) on nitrogen stored in oyster a) tissue and b) shell growth (mg). Initial shell mass (mg) was used as a covariate. We also included the following nested terms as random effects to account for the nested design and non-independence among individuals in Year 1 and 2: Year 1 replicate was nested within Year 1 DO and Year 1 Temp, Year 2 replicate was nested within Year 2 DO and Year 2 Temp, and Year 2 replicate was separately crossed with Year 1 DO, Year 1 Temp, and Year 1 DO x Year 1 Temp and nested within Year 2 DO and Year 2 Temp. Analyses were conducted in R v. 3.6.3 (10) using the lme4 (11), car (12), and emmeans (13) packages.

| | a) N in tissue growth (mg) | | b) N in shell growth (mg) | |
|---|---|---|---|---|
| **Effect** | **F ($df_{num}$, $df_{den}$)** | **p** | **F ($df_{num}$, $df_{den}$)** | **p** |
| Year 1 DO | 3.28 (1,15.5) | 0.09 | 11.12 (1,16.2) | **0.004** |
| Year 1 Temp | 0.17 (1,11.8) | 0.69 | 2.51 (1,14.3) | 0.13 |
| Year 2 DO | 1.22 (1,20.0) | 0.28 | 25.26 (1,20.1) | **<0.0001** |
| Year 2 Temp | 4.42 (1,20.0) | **0.048** | 77.89 (1,20.0) | **<0.0001** |
| Y1 DO*Y1 Temp | 5.68 (1,9.6) | **0.039** | 1.27 (1,7.89) | 0.29 |
| Y1 DO*Y2 DO | 1.88 (1,20.0) | 0.19 | 0.98 (1,19.9) | 0.33 |
| Y1 DO*Y2 Temp | 3.52 (1,20.0) | 0.07 | 12.31 (1,19.8) | **0.002** |
| Y1 Temp*Y2 DO | 0.43 (1,19.9) | 0.52 | 14.40 (1,19.6) | **0.001** |
| Y1 Temp*Y2 Temp | 1.44 (1,19.9) | 0.24 | 0.38 (1,19.7) | 0.55 |
| Y2 DO*Y2 Temp | 5.25 (1,20.0) | **0.03** | 167.46 (1,20.0) | **<0.0001** |
| Y1 Temp*Y1 DO*Y2 DO | 0.38 (1,20.2) | 0.55 | 28.60 (1,20.2) | **<0.0001** |
| Y1 Temp*Y1 DO*Y2 Temp | 9.98 (1,20.1) | **0.005** | 12.34 (1,20.0) | **0.002** |
| Y2 Temp*Y1 DO*Y2 DO | 1.21 (1,20.0) | 0.29 | 11.98 (1,19.9) | **0.002** |
| Y2 Temp*Y1 Temp*Y2 DO | 8.06 (1,19.9) | **0.01** | 5.70 (1,19.7) | **0.027** |
| Y1 Temp*Y1 DO*Y2 Temp*Y2 DO | 0.27 (1,20.1) | 0.61 | 0.27 (1,20.0) | 0.61 |
| Initial Shell Mass (mg) | 720.40 (1,926.0) | **<0.0001** | 1687.2 (1,936.0) | **<0.0001** |

**Table S4.** Mean ± SEM nitrogen (% by weight) for oysters exposed to diel-cycling dissolved oxygen (DO, normoxia/hypoxia) and temperature (ambient/warm) treatments at three-months-old (Year 1) and again one year later (Year 2). See Methods for sample preparation. n = 6 for each treatment combination.

| Year 2 DO | Year 2 Temp | Year 1 DO | Year Temp | Final shell length (mm) | *% total Nitrogen* | | *Nitrogen in growth (mg)* | |
|---|---|---|---|---|---|---|---|---|
| | | | | | Tissue | Shell | Tissue | Shell |
| Normoxia | Ambient | Normoxia | Ambient | 14.01 ± 0.32 | 7.52 ± 0.45 | 0.108 ± 0.008 | 7.29 ± 0.43 | 0.232 ± 0.012 |
| | | | Warm | 13.41 ± 0.35 | 8.82 ± 0.20 | 0.120 ± 0.016 | 7.53 ± 0.70 | 0.248 ± 0.014 |
| | | Hypoxia | Ambient | 13.42 ± 0.36 | 9.57 ± 0.82 | 0.133 ± 0.021 | 8.33 ± 0.49 | 0.274 ± 0.014 |
| | | | Warm | 13.52 ± 0.41 | 8.53 ± 0.28 | 0.100 ± 0.000 | 8.27 ± 0.66 | 0.205 ± 0.011 |
| | Warm | Normoxia | Ambient | 13.97 ± 0.38 | 9.68 ± 1.19 | 0.180 ± 0.016 | 10.37 ± 0.55 | 0.402 ± 0.018 |
| | | | Warm | 13.69 ± 0.40 | 8.62 ± 0.15 | 0.170 ± 0.016 | 8.65 ± 0.55 | 0.372 ± 0.019 |
| | | Hypoxia | Ambient | 13.82 ± 0.40 | 8.60 ± 0.21 | 0.162 ± 0.018 | 9.05 ± 0.62 | 0.354 ± 0.021 |
| | | | Warm | 13.42 ± 0.35 | 8.77 ± 0.24 | 0.130 ± 0.016 | 9.37 ± 0.61 | 0.286 ± 0.015 |
| Hypoxia | Ambient | Normoxia | Ambient | 14.32 ± 0.42 | 9.50 ± 0.99 | 0.148 ± 0.018 | 8.75 ± 0.49 | 0.311 ± 0.015 |
| | | | Warm | 13.74 ± 0.36 | 8.77 ± 0.16 | 0.150 ± 0.022 | 7.78 ± 0.59 | 0.290 ± 0.018 |
| | | Hypoxia | Ambient | 13.76 ± 0.43 | 10.1 ± 1.24 | 0.137 ± 0.020 | 8.39 ± 0.56 | 0.259 ± 0.014 |
| | | | Warm | 13.82 ± 0.37 | 8.67 ± 0.28 | 0.128 ± 0.016 | 8.02 ± 0.54 | 0.260 ± 0.014 |
| | Warm | Normoxia | Ambient | 13.48 ± 0.33 | 8.75 ± 0.19 | 0.150 ± 0.018 | 8.69 ± 0.51 | 0.290 ± 0.013 |
| | | | Warm | 13.02 ± 0.40 | 8.57 ± 0.19 | 0.138 ± 0.020 | 7.82 ± 0.42 | 0.260 ± 0.012 |
| | | Hypoxia | Ambient | 12.87 ± 0.34 | 8.38 ± 0.11 | 0.123 ± 0.016 | 7.37 ± 0.50 | 0.227 ± 0.011 |
| | | | Warm | 13.08 ± 0.33 | 8.93 ± 0.25 | 0.132 ± 0.016 | 8.89 ± 0.56 | 0.258 ± 0.014 |

**Table S5.** Potential nitrogen assimilated by oysters restored to Harris Creek, Maryland based on the final nitrogen content of oysters from the experiment that manipulated early life exposure to diel-cycling dissolved oxygen (DO, normoxic/hypoxic) and warming (ambient/warm) and exposure to hypoxia and warming one year later. Allometric equations were empirically derived from experimental oysters measured at the end of the experiment and, for a subset, also the following spring (n = 269) to extrapolate the tissue and shell masses of restored oysters, which are substantially larger (mean shell length ± SE; 73.6 ± 0.62 mm) than experimental oysters (13.6 ± 0.09 mm). Means and CI were calculated via bootstrapping using the boot R package (14), 1000 iterations, and a normal approximation using oyster shell lengths and densities taken from surveys completed in 2020 of reefs restored in 2013 (121.42 $m^{-2}$, 15).

| Year 2 DO | Year 2 Temp | Year 1 DO | Year 1 Temp | Allometric eq. for tissue mass (mg) from shell length (mm) | $R^2$ | Allometric eq. for shell mass (mg) from shell length (mm) | $R^2$ | n | Mean N in one acre restored reef (kg) | | CI of total N in one acre restored reef |
|---|---|---|---|---|---|---|---|---|---|---|---|
| | | | | | | | | | Tissue | Shell | |
| Normoxia | Ambient | Normoxia | Ambient | ln(tissue mg) = 0.456 + 1.98*ln(shell mm) | 0.79 | ln(shell mg) = 2.158 + 1.63*ln(shell mm) | 0.77 | 73 | 290.34 | 5.09 | 281.9, 309.3 |
| | | | Warm | ln(tissue mg) = 0.985 + 1.76*ln(shell mm) | 0.78 | ln(shell mg) = 2.795 + 1.40*ln(shell mm) | 0.78 | 78 | 224.45 | 3.97 | 219.1, 238.0 |
| | | Hypoxia | Ambient | ln(tissue mg) = 0.487 + 1.94*ln(shell mm) | 0.86 | ln(shell mg) = 2.347 + 1.55*ln(shell mm) | 0.84 | 76 | 320.94 | 5.37 | 311.6, 341.3 |
| | | | Warm | ln(tissue mg) = 0.339 + 1.994*ln(shell mm) | 0.80 | ln(shell mg) = 2.22 + 1.59*ln(shell mm) | 0.79 | 76 | 306.08 | 4.21 | 296.0, 324.9 |
| | Warm | Normoxia | Ambient | ln(tissue mg) = 1.731+ 1.50*ln(shell mm) | 0.74 | ln(shell mg) = 3.343 + 1.20*ln(shell mm) | 0.71 | 75 | 169.76 | 4.36 | 168.1, 180.3 |
| | | | Warm | ln(tissue mg) = 1.851 + 1.44*ln(shell mm) | 0.58 | ln(shell mg) = 3.825 + 1.01*ln(shell mm) | 0.54 | 69 | 131.73 | 2.95 | 130.2, 139.3 |
| | | Hypoxia | Ambient | ln(tissue mg) = 1.487 + 1.55*ln(shell mm) | 0.73 | ln(shell mg) = 3.192 + 1.23*ln(shell mm) | 0.67 | 72 | 146.61 | 3.83 | 145.1, 156.0 |
| | | | Warm | ln(tissue mg) = 1.631 + 1.52*ln(shell mm) | 0.68 | ln(shell mg) = 3.005 + 1.30*ln(shell mm) | 0.71 | 73 | 151.70 | 3.45 | 149.7, 160.7 |
| Hypoxia | Ambient | Normoxia | Ambient | ln(tissue mg) = 1.340 + 1.62*ln(shell mm) | 0.83 | ln(shell mg) = 2.962 + 1.32*ln(shell mm) | 0.82 | 86 | 188.91 | 4.11 | 185.8, 200.4 |
| | | | Warm | ln(tissue mg) = 0.763 + 1.82*ln(shell mm) | 0.80 | ln(shell mg) = 2.533 + 1.47*ln(shell mm) | 0.78 | 88 | 231.33 | 5.16 | 226.5, 246.7 |
| | | Hypoxia | Ambient | ln(tissue mg) = 0.915 + 1.74*ln(shell mm) | 0.84 | ln(shell mg) = 2.573 + 1.44*ln(shell mm) | 0.80 | 78 | 219.97 | 4.30 | 215.2, 233.5 |
| | | | Warm | ln(tissue mg) = 1.436 + 1.55*ln(shell mm) | 0.75 | ln(shell mg) = 2.895 + 1.33*ln(shell mm) | 0.74 | 80 | 140.39 | 3.47 | 138.7, 149.2 |
| | Warm | Normoxia | Ambient | ln(tissue mg) = 0.911 + 1.83*ln(shell mm) | 0.74 | ln(shell mg) = 2.642 + 1.47*ln(shell mm) | 0.72 | 74 | 279.50 | 5.74 | 273.2, 297.6 |
| | | | Warm | ln(tissue mg) = 0.917 + 1.80*ln(shell mm) | 0.83 | ln(shell mg) = 2.586 + 1.47*ln(shell mm) | 0.84 | 81 | 241.98 | 5.00 | 236.7, 257.5 |
| | | Hypoxia | Ambient | ln(tissue mg) = 0.967 + 1.78*ln(shell mm) | 0.74 | ln(shell mg) = 2.755 + 1.41*ln(shell mm) | 0.70 | 74 | 228.88 | 4.09 | 223.4, 242.8 |
| | | | Warm | ln(tissue mg) = 0.765 + 1.86*ln(shell mm) | 0.74 | ln(shell mg) = 2.435 + 1.52*ln(shell mm) | 0.69 | 71 | 280.44 | 5.09 | 273.2, 298.1 |

**2018**

| August | | | | | | |
|---|---|---|---|---|---|---|
| Su | Mo | Tu | We | Th | Fr | Sa |
| | | | 1 | 2 | 3 | 4 |
| 5 | 6 | 7 | **8** | 9 | 10 | 11 |
| 12 | 13 | 14 | 15 | 16 | 17 | 18 |
| 19 | 20 | 21 | 22 | 23 | 24 | **25** |
| 26 | 27 | 28 | 29 | 30 | 31 | |

**2019**

| August | | | | | | |
|---|---|---|---|---|---|---|
| Su | Mo | Tu | We | Th | Fr | Sa |
| | | | | 1 | 2 | 3 |
| 4 | 5 | 6 | 7 | 8 | **9** | 10 |
| 11 | 12 | 13 | 14 | 15 | 16 | 17 |
| 18 | 19 | 20 | 21 | 22 | 23 | 24 |
| 25 | 26 | 27 | **28** | 29 | 30 | 31 |

**Figure S1.** Dates of experiment. The Year 1 exposure began on August 8, 2018 and ran through August 25, 2018. The Year 2 exposure began August 9, 2019 and ran through August 28, 2019. DO cycling days marked in grey (13 days in Year 1, 16 days in Year 2). Temperature treatments were applied each day of the experiment (18 days in Year 1, 20 days in Year 2)

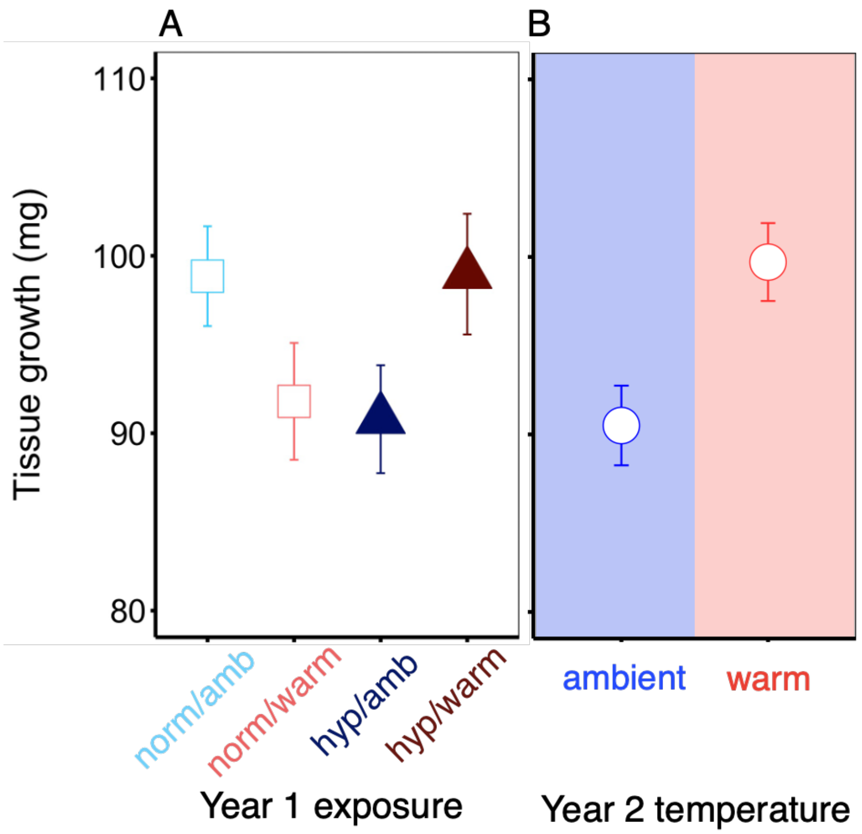

**Figure S2.** Tissue growth (mg) of oysters exposed to (A) diel-cycling dissolved oxygen (normoxia/hypoxia) and temperature (ambient/warm) treatments at three-months-old (Year 1) and (B) different temperatures (ambient/warm) one year later (Year 2). Symbol colors indicate different Year 1 treatment combinations and shaded background colors indicate different Year 2 temperatures. Data are means ± SEM.

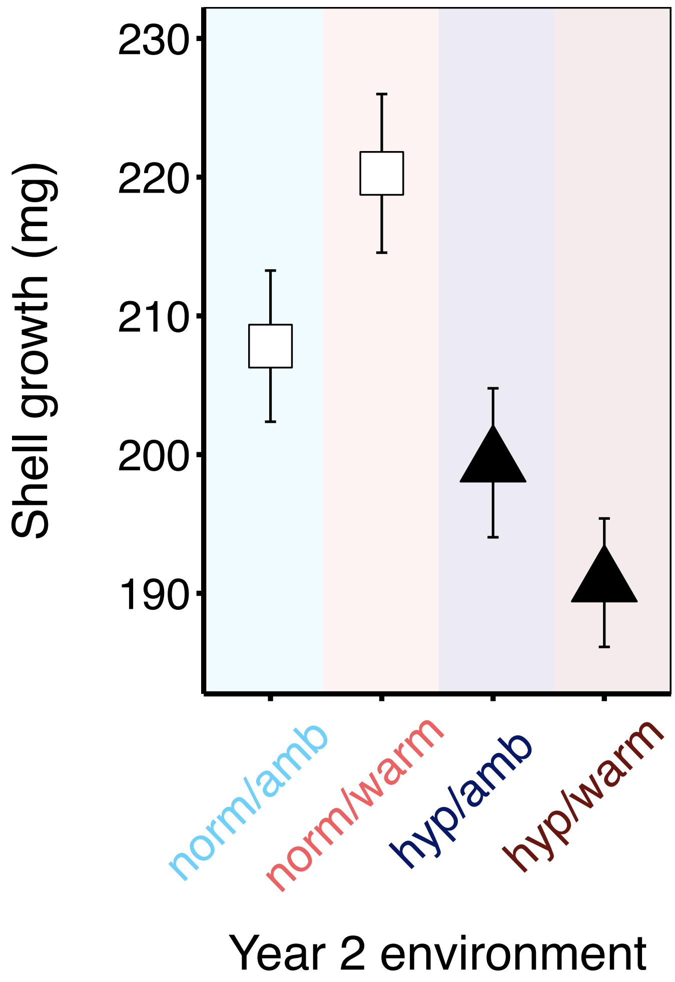

**Figure S3.** Shell growth (mg) of oysters exposed to diel-cycling dissolved oxygen (normoxia/hypoxia) and temperature (ambient/warm) treatments in Year 2 of the experiment. Data are means ± SEM.

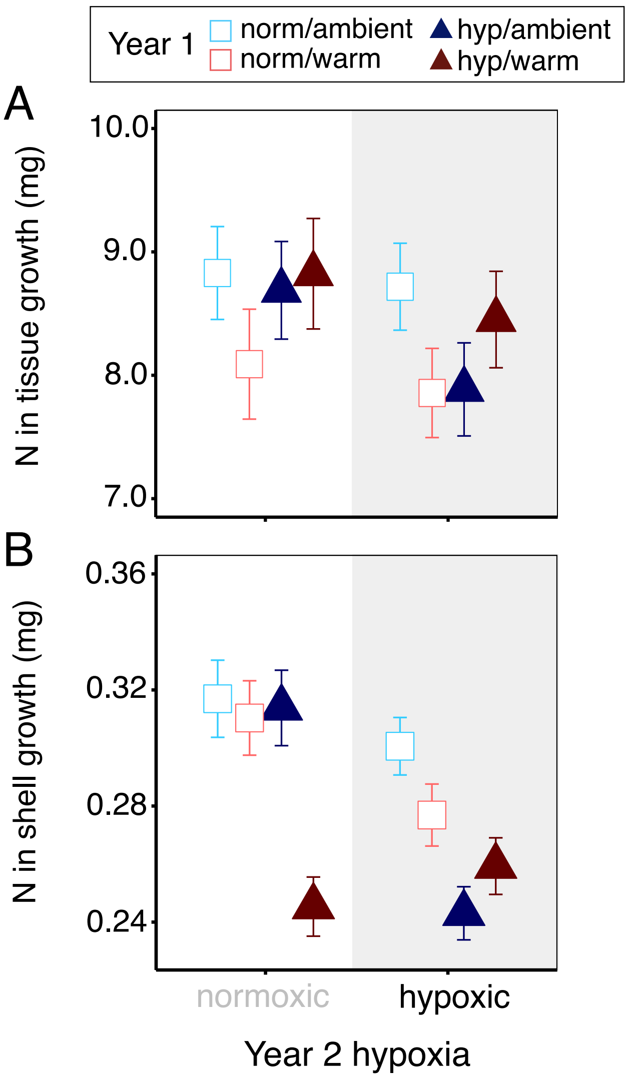

**Figure S4.** Nitrogen bioassimilated into oyster (A) tissue and (B) shell growth (mg) at different Year 2 diel-cycling DO treatments (x-axis) for oysters exposed to diel-cycling dissolved oxygen (normoxia/hypoxia) and temperature (ambient/warm) treatments at three-months-old (Year 1). Empirically derived percent nitrogen by weight provided for each treatment combination in Table S4. Data are means ± SEM.
